# Replication Fork Uncoupling Causes Nascent Strand Degradation and Fork Reversal

**DOI:** 10.1101/2021.10.13.464136

**Authors:** Tamar Kavlashvili, James M Dewar

## Abstract

Genotoxins cause nascent strand degradation (NSD) and fork reversal during DNA replication. NSD and fork reversal are crucial for genome stability and exploited by chemotherapeutic approaches. However, it is unclear how NSD and fork reversal are triggered. Additionally, the fate of the replicative helicase during these processes is unknown. We developed a biochemical approach to study synchronous, localized NSD and fork reversal using *Xenopus* egg extracts. We show that replication fork uncoupling stimulates NSD of both nascent strands and progressive conversion of uncoupled forks to reversed forks. The replicative helicase remains bound during NSD and fork reversal, indicating that both processes take place behind the helicase. Unexpectedly, NSD occurs before and after fork reversal, indicating that multiple degradation steps take place. Overall, our data show that uncoupling causes NSD and fork reversal and identify key steps involved in these processes.

## Introduction

During DNA synthesis genotoxins can cause replication forks to stall. This leads to regression of the replication fork junction and reannealing of the parental DNA strands to generate a reversed fork (Supplemental Fig. S1A,i-ii, fork reversal)^1–4^, which facilitates bypass of DNA lesions and allows for global control of replication fork progression^5, 6^. Fork reversal enzymes can also restore reversed forks, resulting in interconversion between replication forks and reversed forks (Supplemental Fig. S1A,ii-i)^1, 6, 7^. Stalling also leads to degradation of both nascent DNA strands by nucleases (nascent strand degradation, NSD)^8–10^. NSD is thought to target the regressed arm of a reversed fork (Supplemental Fig. S1A,ii-iii) to convert the reversed fork back to a replication fork so that DNA replication can restart (Supplemental Fig. S1A,iii-iv)^1–4^. NSD can be inferred as an initial response to stalling based on the roles of nucleases in restarting DNA synthesis^8, 11^. However, NSD can only be observed after prolonged treatment with genotoxic agents^8, 12^ or inactivation of negative regulators^9, 10, 13^ and has not formally been demonstrated to be an initial response. Additionally, many questions about the mechanism of NSD and fork reversal remain, and it is unclear how these processes are triggered.

Fork reversal and NSD can be triggered by a variety of genotoxic agents^8, 14–19^. Hydroxyurea, which stimulates both fork reversal and NSD^8, 9, 13, 14^, stalls replication forks indirectly by inhibiting ribonucleotide reductase and depleting cellular nucleotides, resulting in DNA polymerase stalling^20^. The replicative helicase continues to unwind, even in the absence of DNA polymerase activity, resulting in replication fork uncoupling^21, 22^. Uncoupling correlates well with fork reversal^14^, suggesting that uncoupling may cause fork reversal and possibly NSD. However, once the replicative helicase uncouples, it effectively stalls^23, 24^ raising the possibility that helicase stalling causes fork reversal and NSD. Accordingly, helicase stalling lesions such as topoisomerase poisons and crosslinking agents can also cause fork reversal^14–17^ and NSD^8, 18, 25^. Moreover, recent data suggests that neither replication fork uncoupling nor stalling of the replicative helicase is sufficient to trigger NSD unless combined with reactive oxygen species^26^. Thus, it is unclear whether fork stalling leads to NSD and fork reversal by causing replication fork uncoupling, helicase stalling, or some other event.

Many proteins are now implicated in fork reversal and NSD^1–4^. The helicases SMARCAL1, HLTF, ZRANB3, and FBH1 are important for regression of the fork junction^27–30^. The recombination proteins RAD51 and RAD52 are also important for fork reversal^8, 10, 14, 31^, though it is unclear exactly how. The exonucleases DNA2 and MRE11 as well as many others are required for degradation^1–3^. NSD is also negatively regulated by ‘fork protection’ factors that prevent excessive NSD from occurring^9, 13^, which would otherwise result in genome instability^9, 11, 28^. The prototypical fork protection protein BRCA2 works by stabilizing RAD51 on regressed arms^9, 11, 31^, but in most cases the exact mechanism by which NSD is inhibited by fork protection factors is unclear. Furthermore, fork reversal, NSD, and fork protection all require multiple proteins acting in a non-redundant manner and the mechanistic basis for this is unclear^1–4^. Thus, although we know many proteins that participate in fork reversal, NSD and fork protection, their exact function is unclear. This is important to address because excessive NSD contributes to the cytotoxicity of certain chemotherapeutics and amelioration of NSD can cause chemoresistance^28, 32, 33^.

Key mechanistic questions about fork reversal and NSD remain. First, the fate of the replicative helicase is unclear. Strand annealing in the vicinity of the replicative helicase was recently reported to promote its transition from single-stranded to double-stranded DNA^34^, which would be expected to trigger replicative helicase unloading during fork reversal^35^. Accordingly, unloading of the replicative helicase is required for fork reversal during one of the pathways for DNA interstrand cross-link repair^17^. However, the replicative helicase can be detected at replication forks in response to hydroxyurea treatment^36^, which would be expected to cause fork reversal^14^ and NSD^8^. This raises the possibility that the replicative helicase remains present during fork reversal and NSD. Second, it is unclear which DNA structures are targeted for degradation. Current models propose that degradation begins at reversed forks^1–4^, consistent with a requirement for fork reversal during extensive NSD^31^. However, it is unclear whether degradation continues at replication forks once the regressed arm is lost and whether the same is true during the more limited NSD that occurs during restart of DNA synthesis^8^.

Conventional approaches lack the temporal and spatial resolution as well as the sensitivity needed to address the questions outlined above. To overcome these limitations, we used *Xenopus* egg extracts to develop a highly sensitive, synchronous, and locus-specific approach to study NSD and fork reversal. NSD induced by this approach involves DNA2, as previously reported. NSD degrades both nascent strands and replication forks are progressively converted to reversed forks. Using this approach, we show that replication fork uncoupling, rather than helicase stalling, elicits both fork reversal and nascent strand degradation. Additionally, the replicative helicase remains bound throughout both fork reversal and NSD. Since replisome removal pathways are not activated, this observation strongly suggests that the replisome remains on single-stranded DNA, while fork reversal and NSD take place behind the uncoupled helicase. Furthermore, our data reveal that degradation occurs at replication forks before and after reversal, indicating that two degradation steps occur, one at uncoupled forks and one at reversed forks. Overall, our research identifies a trigger for NSD and reveals key mechanistic insights into NSD and fork reversal.

## Results

### NSD is an initial response to fork stalling

We first tested whether NSD is an initial response to fork stalling. To this end, we examined the stability of nascent DNA strands following treatment with aphidicolin, during DNA replication in *Xenopus* egg extracts. Aphidicolin is a DNA polymerase inhibitor that mimics the effects of hydroxyurea (HU)^14, 21, 22, 37^, which is commonly used to induce NSD in cells^8, 9, 13, 38^, and can be used instead of HU^39^ to induce fork reversal and NSD in *Xenopus* egg extracts^27^. The *Xenopus* system we employed^40^ is highly sensitive to even small changes in nascent DNA signal^41^ which afforded us the opportunity to monitor even low levels of degradation. DNA replication was initiated on plasmid templates that replicate semi-synchronously from a single origin per plasmid^42^ and nascent DNA strands were labeled by inclusion of radiolabeled nucleotides in the reaction (Supplemental Fig. S1B). Shortly after initiation, DNA synthesis was stalled by addition of aphidicolin (Supplemental Fig. S1B). Samples were withdrawn at different time points, purified, and digested so that total radioactive signal could be quantified relative to a loading control (Supplemental Fig. S1C). In vehicle-treated extracts, radioactive signal increased and then plateaued (Supplemental Fig. S1D, lanes 1-5; Supplemental Fig. S1E-F), indicating that a single round of replication was completed. In contrast, in aphidicolin-treated extracts, signal was reduced by 60 minutes and mostly gone by 120 minutes (Supplemental Fig. S1D, lanes 6-10; Supplemental Fig. S1E-F), indicating that degradation took place. To test whether the degradation corresponded to both strands or nascent strands only, we radiolabeled parental strands and repeated the same experiment (Supplemental Fig. S1G). Parental strand signal was relatively stable and there was little difference between aphidicolin and vehicle treatment (Supplemental Fig. S1H, compares lanes 1-5 and 6-10; Supplemental Fig. S1I-J). Moreover, there was no appreciable loss of parental strand signal for the first 120 minutes (Supplemental Fig. S1I-J), during which time most of the nascent strands were degraded (Supplemental Fig. S1E-F). Thus, most signal loss was due to degradation of nascent DNA strands. Our data show that NSD occurs shortly after fork stalling and thus is an initial response, as expected based on cellular studies^8, 11^.

### A synchronous biochemical approach to study NSD

Our initial approach (Supplemental Fig. S1B-C) stalled replication forks shortly after initiation of DNA synthesis and resulted in a heterogenous population of replication forks that were replicated to different extents and at different locations (Supplemental Fig. S1D, lane 6). To carefully monitor NSD with high temporal and spatial resolution, we localized forks to a *lac* repressor (LacR) barrier^41^ before NSD was induced (Fig. 1A). Plasmid DNA harboring a *lacO* array was pre-incubated with LacR and then replicated in *Xenopus* egg extracts, which gave rise to θ structures (Fig. 1A; Fig. 1B, lane 1), as expected^41^. Forks were then stalled with aphidicolin (Fig. 1A) and IPTG was simultaneously added to disrupt the LacR array to allow uncoupling of the replicative helicase, which also occurs following HU treatment in cells^21, 22^. Upon aphidicolin and IPTG treatment, θs were converted to θ* structures (Fig. 1B, lanes 2-5), which arose from reannealing of unwound DNA strands in the detergent-treated samples (Supplemental Fig. S2A,B) and showed that uncoupling occurred. To remove any topological effects, we purified the DNA and performed restriction digests to visualize the replication fork structures (Fig. 1D)^41^. Double-Ys disappeared over time without any appreciable increase in linear products of replication (Fig. 1E, lanes 2-5; Fig. 1F; Supplemental Fig. S2C). Thus, loss of the Double-Y species was due to degradation of nascent DNA strands. Accordingly, the majority of nascent strand DNA signal was lost (Supplemental Fig. S2D). These results show that we have developed a synchronous biochemical approach to induce NSD.

**Figure 1:**
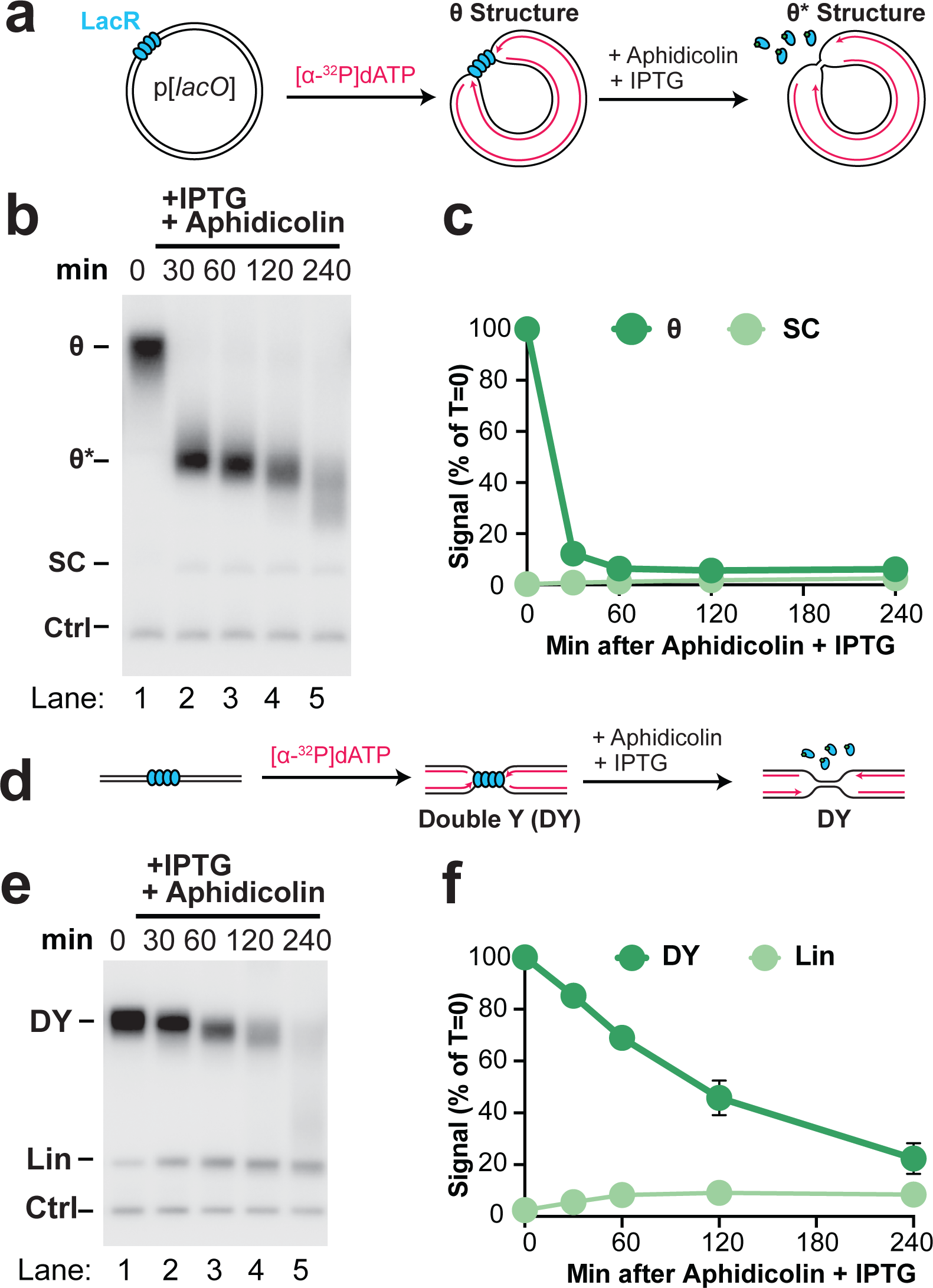
Induction of synchronous and localized nascent strand degradation *in vitro*. *(A)* Plasmid DNA harboring a LacR array was replicated using *Xenopus* egg extracts and dATP[α-^32^P] was added to label newly-synthesized DNA strands. Once forks were localized at the LacR barrier replication was restarted by addition of IPTG to remove the barrier and aphidicolin was added to stall replication forks. *(B)* Samples from (a) were separated on an agarose gel and visualized by autoradiography. As a loading control (Ctrl) the reactions include a fully-replicated plasmid. See also Supplemental Fig. S2A-B. *(C)* Quantification of θ structures and supercoiled monomers (SC) from (b). Mean ± S.D., n=3 independent experiments. *(D)* Samples from (a) were digested with XmnI to allow unambiguous identification of replication fork structures (DYs) and linear products of replication (Lin). See Supplemental Fig. S2A,i for a restriction map. *(E)* Samples from (d) were separated on an agarose gel and visualized by autoradiography. *(F)* Quantification of DY and Lin structures in (e). Mean ± S.D., n=5 independent experiments. See also Supplemental Fig. S2C-D.

### NSD involves the DNA2 exonuclease

In other settings NSD involves the DNA2 or MRE11 exonucleases (Supplemental Fig. S3A)^2, 8, 9, 30^. To determine which nuclease was involved in the NSD we observed, we used the small molecule inhibitors C5 (DNA2-i)^43^ and Mirin (MRE11-i)^44^ to inactivate these nucleases in *Xenopus* egg extracts. To determine the effectiveness of these inhibitors, we first targeted DNA2 and MRE11 during Double-Strand Break (DSB) resection, which involves both enzymes in *Xenopus* egg extracts (Supplemental Fig. S3A) and is blocked by inactivation of both DNA2 and MRE11^45^. We incubated radiolabeled linear DNA substrates in extracts and monitored degradation, which serves as a read-out for resection (Fig. 2A). In the vehicle control the linear substrate was rapidly degraded (Fig. 2B, lanes 1-3; Fig. 2C). We also observed a small number of additional products that arose from end-joining (Fig. 2B, EJ(L) and EJ(C)). As expected, DNA2-i and MRE11-i each inhibited resection (Fig. 2B, lanes 4-6, 6-9, Fig. 2C, lanes 1-3), while a combination of both inhibitors almost completely blocked resection (Fig. 2B, lanes 10-12, Fig. 2C). This demonstrated that both DNA2-i and MRE11-i were able to efficiently inhibit their targets.

**Figure 2:**
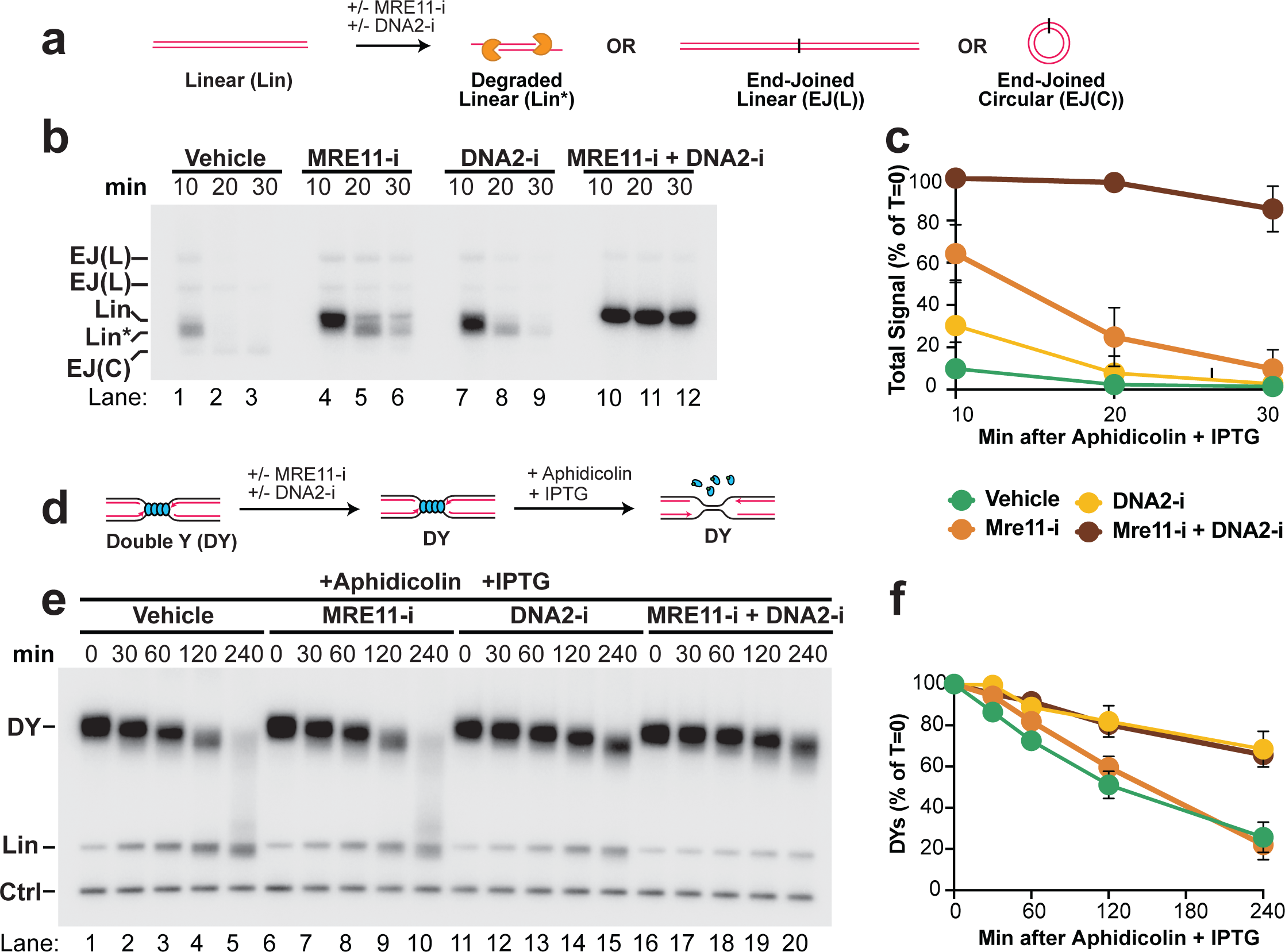
Nascent strand degradation involves DNA2 but not MRE11. *(A)* Linear radiolabeled DNA was incubated in *Xenopus* egg extracts in the presence of MRE11 inhibitor (MRE11-i) or DNA2 inhibitor (DNA2-i) or both. *(B)* Samples from (a) were separated on an agarose gel and visualized by autoradiography. *(C)* Quantification of total signal from (b). Mean ± S.D., n=3 independent experiments. *(D)* Forks were localized to a LacR barrier and NSD was induced by addition of IPTG and aphidicolin in the absence or presence of MRE11-i and DNA2-i. Purified DNA was subjected to restriction digest so that replication fork structures could be visualized as DYs. *(E)* Samples from (d) were separated on an agarose gel and visualized by autoradiography. See also Supplemental Fig. S3C-D. *(F)* Quantification of DY signal from (e). Mean ± S.D., n=3 independent experiments. See also Supplemental Fig. S3E-F.

To determine the roles of DNA2 and MRE11 in our system, we induced NSD (as in Fig. 1) in the presence of either MRE11-i, DNA2-i or both (Fig. 2D). In vehicle treated extracts, we again observed efficient NSD (Fig. 2E, lanes 1-5; Fig. 2F). Interestingly, DNA2-i treatment blocked the majority of NSD (Fig. 2E, lanes 1-5,11-15; Fig. 2F) while MRE11-i had little effect either alone (Fig. 2E, lanes 1-10; Fig. 2F) or in combination with DNA2-i (Fig. 2E, lanes 1-5 and 16-20; Fig. 2F). Importantly, neither MRE11-i nor DNA2-i prevented uncoupling (Supplemental Fig. S3C-D). These data show that the NSD we observe involves DNA2 but not MRE11, consistent with previous reports^8–10, 13^ (but see also^33, 38^).

### NSD involves fork reversal

NSD typically involves replication fork reversal ^8, 13, 31^. To test whether this was also the case in our experiments, we performed 2D gel electrophoresis to monitor the DNA structures formed during NSD (Fig. 3A,B). At early time points, signal was broadly distributed throughout the bubble and double-Y arcs, due to initiation of replication throughout the plasmid (Fig. 3C,i). Immediately prior to induction of NSD, most signal was present as a discrete spot on the double-Y arc, corresponding to forks localized to the LacR barrier (Fig. 3C,ii, DYs). Interestingly, we also observed a small population of replication forks that did not represent canonical replication intermediates and were therefore altered replication fork structures i.e. remodeled forks (Fig. 3C,ii, RFs). During NSD, the abundance of Double-Ys declined (Fig. 3C,ii-iv; Fig. 3D) while remodeled forks increased ∼3 fold (Fig. 3C,ii-iv, Fig. 3E), indicating that Double-Ys were converted to remodeled forks. Importantly, the frequency of reversed forks increased over time indicating progressive conversion of Double-Ys to these species (Fig. 3E). Moreover, the mobility of both Double-Ys and remodeled forks decreased over time (Fig. 3C,ii-iv) indicating that both types of structures included degraded molecules. Thus, nascent strand degradation progressive remodeling of replication forks. However, it is unclear from this data whether the remodeled forks correspond to reversed forks or other DNA structures.

**Figure 3:**
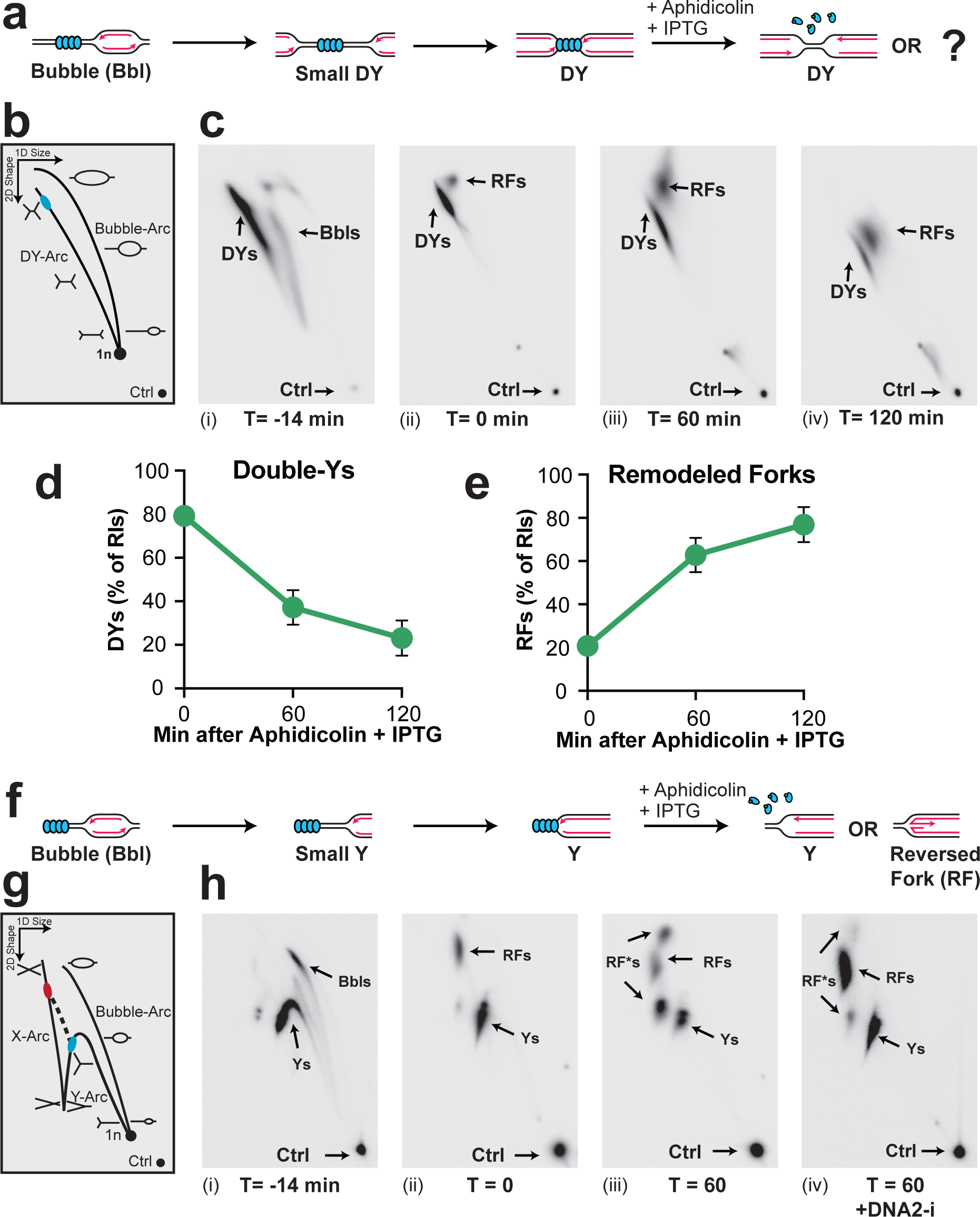
Nascent strand degradation involves fork reversal. *(A)* Cartoon depicting the different DNA structures formed during replication of plasmid DNA harboring a LacR array and induction of NSD. The structures depicted arise from restriction digest using an enzyme that cuts the plasmid template once. *(B)* 2-D gel migration pattern of the DNA structures depicted in (a). The blue spot on the Double Y-arc (DY-arc) indicates the expected migration of DY structures localized to the LacR array. *(C)* DNA structures from (a) were analyzed by 2-D gel electrophoresis and visualized by autoradiography. T=0 is the point at which IPTG and aphidicolin were added. DY, bubble (Bbl), and remodeled fork (RF) structures are indicated. *(D)* Quantification of DY structures from (c) as a percentage of replication intermediates (RIs) at each time point. Mean ± S.D., n=3 independent experiments. See also Supplemental Fig. S4A. *(E)* Quantification of RF structures from (c) as a percentage of RIs at each time point. Mean ± S.D., n=3 independent experiments. See also Supplemental Fig. S4B. *(F)* Cartoon depicting the same structures in (a) after restriction digest using two different enzymes to release a single large fragment. See also Supplemental Fig. S4J. *(G)* 2-D gel migration pattern of the DNA structures depicted in (f). The blue spot on the Y-arc indicates the expected migration of Y structures localized to the LacR array. The red spot indicates the expected migration of reversed fork structures arising from the Y structures within the black spot. *(H)* DNA structures from (f) were analyzed by 2-D gel electrophoresis and visualized by autoradiography. T=0 is the point at which IPTG and aphidicolin were added.

Migration of the remodeled forks (Fig. 3C) did not correspond to canonical Holliday junctions or hemicatenanes (Supplemental Fig. S4C,i-ii, D,i-ii) but was potentially consistent with fork reversal (Supplemental Fig. S4C,iii, D,iii). However, the DNA structures resulting from reversal of one or both forks of a double-Y molecule are undefined so a clear assignment could not be made. Furthermore, the remodeled forks could, in principle, arise from hemicatenane or Holliday junction formation in the wake of one or both forks (Supplemental Fig. S4C,iv,v, D,iv,v). To further examine the identity of the remodeled forks, we treated the double-Y and remodeled fork species with RuvC, which cleaves reversed forks and Holliday junctions, but not hemicatenanes or other replication fork structures^17^. RuvC treatment had essentially no effect on Double-Y structures (Supplemental Fig. S4E-G), as expected, but reduced the abundance of remodeled forks by ∼5-fold (Supplemental Fig. S4E-G). This shows that the remodeled forks contain four-way junctions and thus correspond to either reversed forks (Supplemental Fig. S4H,i) or replication forks followed by a Holliday junctions i.e. D-loops (Supplemental Fig. S4H,ii).

The migration of reversed forks and D-loops during 2-D gel electrophoresis is well defined for DNA molecules that contain only a single fork structure (Supplemental Fig. S4H-I)^46, 47^. We therefore treated NSD intermediates with a pair of restriction enzymes to yield two replication forks of differing sizes that could be separated by gel electrophoresis (Supplemental Fig. S4J). This allowed us to perform 2-D gel electrophoresis of the larger fork only (Fig. 3F) to determine whether the remodeled forks corresponded to reversed forks (Fig. 3G). Shortly after initiation, signal was distributed between bubble and Y-arcs, due to initiation of replication throughout the plasmid (Fig. 3H,i). Immediately prior to NSD, most signal was present as a discrete spot on the Y arc, corresponding to forks localized to the LacR barrier (Fig. 3H,ii, Ys). A small population of remodeled forks was observed immediately prior to NSD (Fig. 3H,ii). Importantly, this species migrated mid-way down the X spike (Fig. 3H,ii, RFs) in the expected position for a reversed fork (Supplemental Fig. S4I,i) and not along the Y-arc, as would be expected for a D-loop (Supplemental Fig. S4I,ii). Surprisingly, this species did not increase in abundance during NSD (Fig. 3H,iii, RFs) in contrast to the remodeled forks observed previously (Fig. 3C). Instead, we observed two additional species during NSD that also migrated along the X-spike with a similar mobility to reversed forks (Fig. 3H,iii, RF*s). These species almost completely disappeared following DNA2-i treatment (Fig. 3H,iv), which also led to a corresponding increase in RF signal (Fig. 3H,iv). Thus, during NSD reversed forks are generated that are normally degraded by DNA2. Overall, these data show that NSD induced by our approach involves progressive conversion of replication forks to reversed forks, which are then degraded by DNA2.

### Uncoupling elicits NSD and fork reversal

NSD in cells involves degradation of both DNA strands^8, 9, 13^. To test whether this was the case in our system, DNA intermediates were purified, restriction digested, then separated on an agarose gel under denaturing conditions (Fig. 4A). This approach allowed us to monitor the loss of intact nascent strands (Fig. 4A, LWS) as a more direct read-out for the onset of NSD than measurement of Double-Y signal (as in Fig. 1; Fig. 2). Upon addition of IPTG and aphidicolin, intact strands were readily lost (Fig. 4B, LWS; Fig. 4C) and degraded strands could be visualized (Fig. 4B, lane 2, Deg). The disappearance of essentially all intact strands (Fig. 4B, lanes 1-4; Fig. 4c) indicated that both leading and lagging strands were degraded. Further analysis of both sets of nascent strands (Fig. 4A, RWS and LWS) confirmed that both strands were degraded (Supplemental Fig. S5A-C). Thus, the NSD we observe involves degradation of both nascent strands.

**Figure 4:**
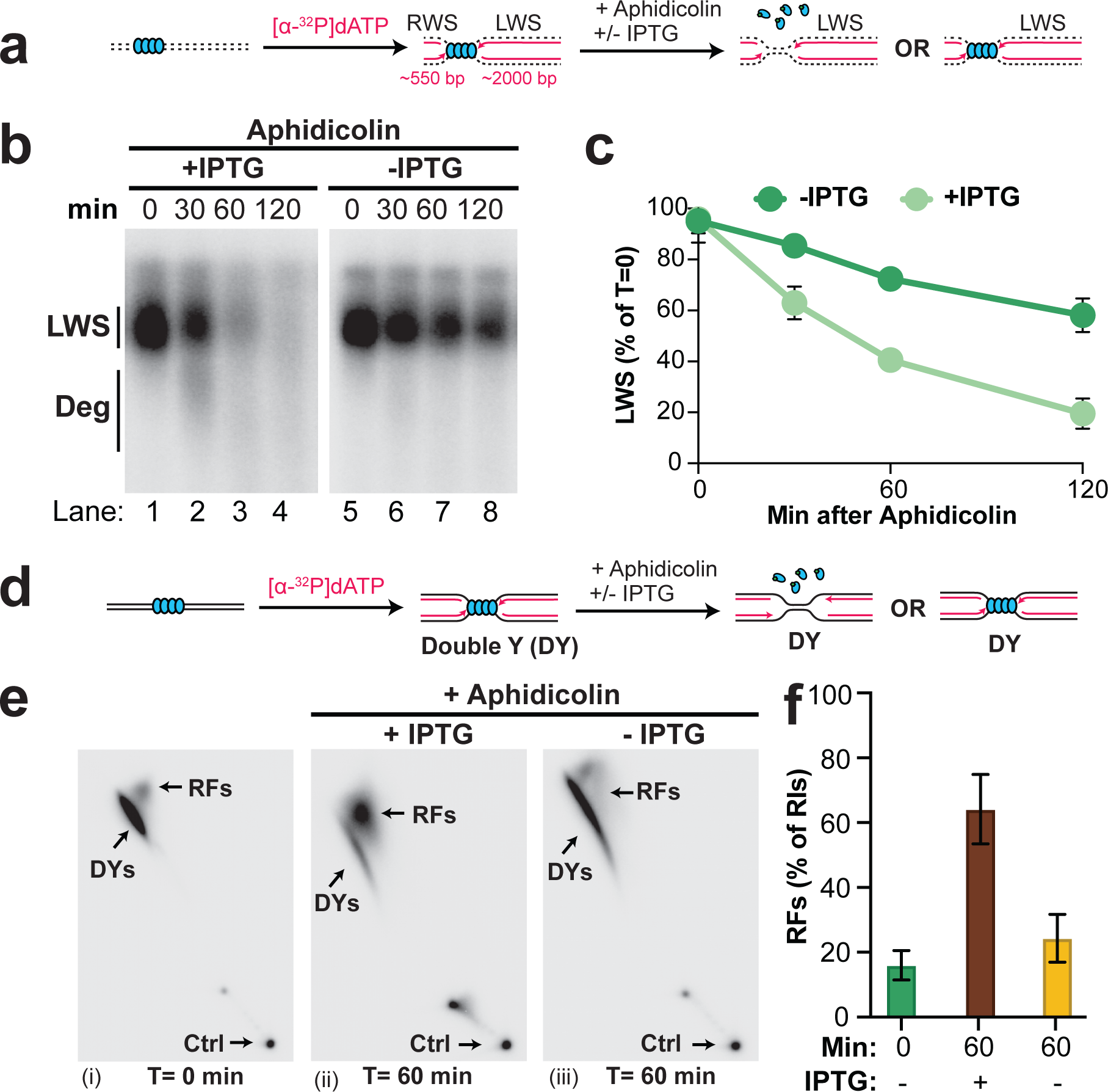
Uncoupling promotes nascent strand degradation and fork reversal. *(A)* Forks were localized to a LacR barrier and aphidicolin was added in the absence or presence of IPTG. Purified DNA was restriction digested to yield leftward strands (LWS) and rightward strands of different sizes. *(B)* DNA structures from (a) were separated on an alkaline denaturing agarose gel and visualized by autoradiography. Intact and degraded (deg) LWS are indicted. See also Supplemental Fig. S5B-C,E-F. *(C)* Quantification of intact LWS strands from (b). Mean ± S.D., n=3 independent experiments. *(D)* Samples from (a) were subjected to restriction digest so that replication fork structures could be visualized as DYs. *(E)* Double-Y (DY) and reversed fork (RF) structures from (d) were separated by 2-D gel electrophoresis (as in Fig. 3B-C). *(F)* Quantification of RFs from (e) as a percentage of replication intermediates (RIs) at each time point. Mean ± S.D., n=3 independent experiments. See also Supplemental Fig. S5J.

Fork stalling by aphidicolin results in replication fork uncoupling^48^ and helicase stalling^23^. We therefore wanted to test which of these events trigger NSD and fork reversal. To directly test this, we localized forks to a LacR barrier, added aphidicolin, and either added or omitted IPTG to facilitate or inhibit helicase uncoupling, respectively (Supplemental Fig. S5D-E). Addition of IPTG resulted in uncoupling (Supplemental Fig. S5E-F) and NSD (Fig. 4A-C), as described above. When IPTG was omitted, uncoupling was strongly inhibited (Supplemental Fig. S5E-F) but not blocked, as expected^41^. Importantly, omission of IPTG caused intact strands to persist (Fig. 4B, lanes 5-8; Fig. 4C). Omission of IPTG also caused double-Ys to persist when total degradation was monitored (as in Fig. 1; Fig. 2), in the absence or presence of DNA2-i (Supplemental Fig. S5G-I). These data show that replication fork uncoupling, independent of stalling, causes NSD.

We next tested whether fork reversal is also stimulated by replication fork uncoupling. To this end, we performed 2-D gel analysis to monitor the abundance of reversed forks during NSD (as in Fig. 3) in the absence or presence of IPTG (Fig. 4D). Reversed forks accumulated ∼3-fold in the presence of IPTG (Fig. 4E,i-ii; Fig. 4F) as previously shown (Fig. 3E). Strikingly, omission of IPTG essentially blocked the accumulation of reversed forks (Fig. 4,E,iii; Fig. 4F). Thus, replication fork uncoupling induces fork reversal. Overall, our data definitively show that replication fork uncoupling can stimulate both NSD and fork reversal.

### The CMG helicase is retained during NSD and fork reversal

The fate of the CMG helicase during fork reversal and nascent strand degradation is largely elusive. One model^1^ (Supplemental Fig. S6A,i) is that CMG translocates onto double-stranded DNA (dsDNA) during fork reversal^34^, which would lead to replisome unloading^35^. Accordingly, CMG unloading has been reported to be required for fork reversal^17^. Another possibility we considered (Supplemental Fig. S6A,ii) is that the replisome remains associated with the nascent DNA, consistent with long-term retention of the replisome^36^ under conditions known to cause fork reversal^14^. To determine the fate of the replisome during NSD and fork reversal, we induced NSD then recovered chromatin at different time points and performed Western blotting to monitor the abundance of replisome components (Fig. 5A). As expected, the single-stranded DNA binding protein RPA accumulated and then plateaued (Fig. 5B; Supplemental Fig. S6B), consistent with the presence of high levels of ssDNA from both uncoupling and NSD. Importantly, levels of the CMG components MCM6 and CDC45 were essentially unaltered for the first 60 minutes (Fig. 5B, lanes 1-3; Fig. 5C) despite most NSD and fork reversal taking place within this time frame (Fig. 3; Fig. 4). Thus, our data shows that the CMG helicase remains bound to DNA during NSD and fork reversal.

**Figure 5:**
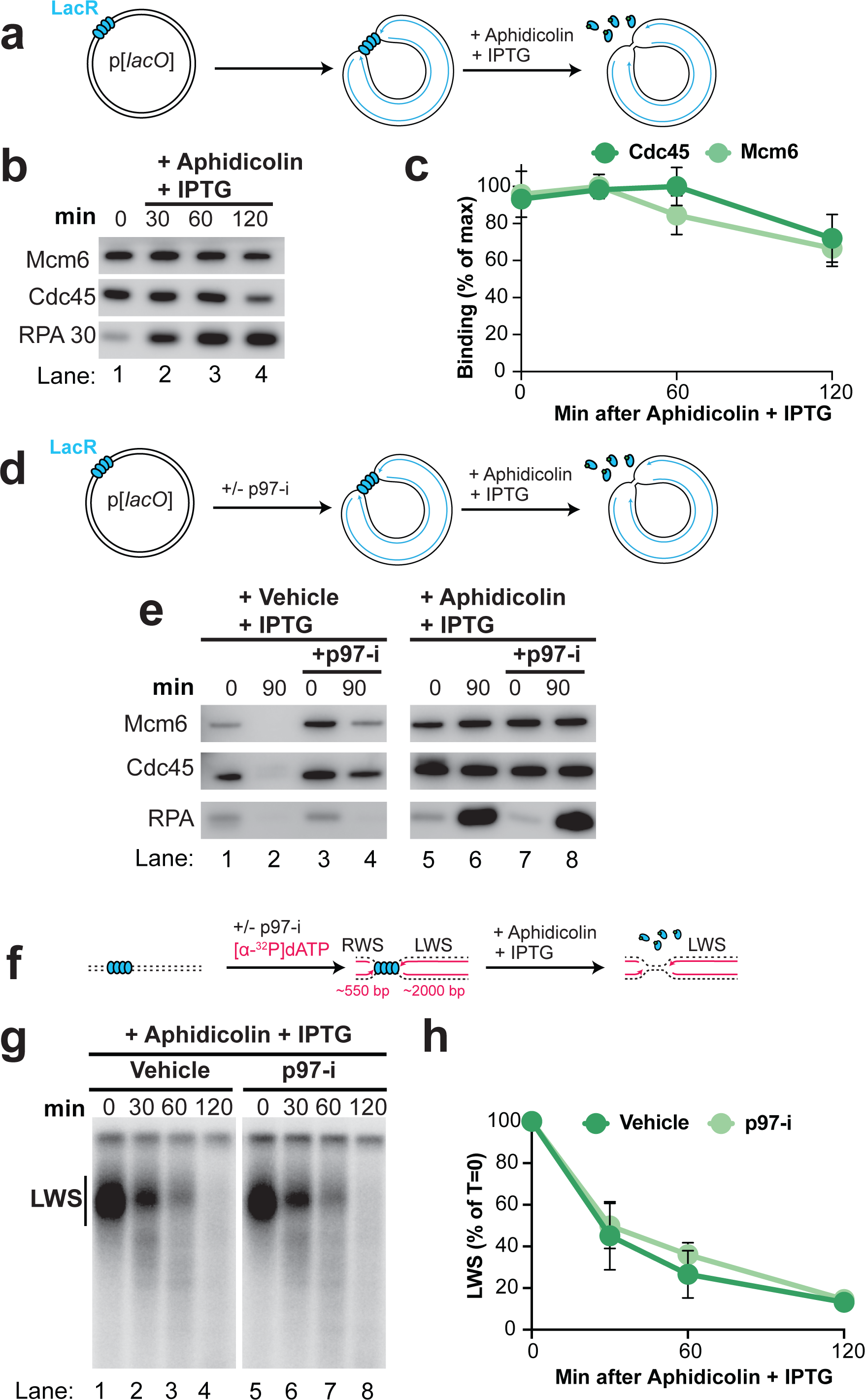
The replisome remains associated with DNA during nascent strand degradation. *(A)* Forks were localized to a LacR barrier and NSD was induced by addition of IPTG and aphidicolin. At different time points chromatin-bound proteins were recovered. *(B)* Proteins from (awere detected by Western Blotting. *(C)* Quantification of CDC45 and MCM6 from (b). Mean ± S.D., n=3 independent experiments. See also Supplemental Fig. S6B. *(D)* NSD was monitored as in (a) in the absence or presence of p97-i. *(E)* Proteins from (d) were detected by western blotting. *(F)* NSD was induced as in (d) and dATP[α-^32^P] was included to radiolabel nascent DNA strands. Purified DNA was restriction digested to yield leftward strands (LWS) and rightward strands of different sizes. *(G)* Samples from (f) were separated on an alkaline denaturing agarose gel and visualized by autoradiography. *(H)* Quantification of LWS from (g). Mean ± S.D., n=3 independent experiments.

Although replisome components were relatively stable at early time points during our experiments, we noticed an ∼25% decrease in MCM6 and CDC45 signal by 120 minutes (Fig. 5B, lanes 3-4; Fig. 5C). While this was not enough to account for all NSD events (Fig. 4C), we wondered whether a subset might require replisome unloading. To test this, we monitored the onset of NSD in the presence of a small molecule inhibitor of p97 (p97-i), which inhibits all known replisome unloading pathways ^41, 49^ (Fig. 5D). When replication forks were allowed to complete DNA synthesis in the absence of aphidicolin, p97-i led to retention of MCM6 and CDC45, but not RPA (Fig. 5E, lanes 1-4), consistent with a specific role for p97 in unloading the replicative helicase and not other replication fork components^42^. When aphidicolin was added, p97-i had little effect on levels of CDC45 or MCM6 (Fig. 5E, lanes 5-8) consistent with replisomes being largely stable at early time points. Accordingly, we monitored the onset of NSD in presence of p97-i (Fig. 5F, as in Fig. 4) and observed no measurable difference in degradation compared to the control (Fig. 5G-H). These data rule out any requirement for replisome unloading in the vast majority of NSD events we observe.

### NSD can occur before fork reversal

Fork reversal is thought to precede NSD^13, 27–29, 31^ (Supplemental Fig. S7A,i). We were therefore surprised by two observations that were potentially with this model. First, 2-D gel analysis showed that the mobility of Double-Ys decreased over time, indicating that these molecules were degraded (Fig. 3C,ii-iv). In principle this could be explained by degradation of reversed forks and subsequent conversion back to Y-shaped forks, as proposed by current models (Supplemental Fig. S7A,i). Second, under conditions that did not induce fork reversal (Fig. 4E,i,iii; Fig. 4F) degradation by DNA2 still occurred (Supplemental Fig. S5H, lanes 1-10; S5I). These data suggest that replication forks can be degraded prior to fork reversal (Supplemental Fig. S7A,iii).

To test whether degradation of Y-shaped forks (double-Ys) arose from degradation of reversed forks, as proposed by current models (Supplemental Fig. S7A,i), or degradation prior to fork reversal, as suggested by our data (Supplemental Fig. S7A,iii) we took advantage of our ability to inhibit degradation using DNA2-i (as in Fig. 2). If degradation begins at reversed forks and converts them to replication forks (Supplemental Fig. S7A,i) then DNA2-i treatment should lead to a decrease in Y-shaped forks (Supplemental Fig. S7A,ii). Alternatively, if degradation begins at Y-shaped forks (Supplemental Fig. S7A,iii), as suggested by our data (above), then DNA2-i treatment should lead to an accumulation of Y-shaped forks (Supplemental Fig. S7A,iv). To distinguish these possibilities, we induced NSD in the presence or absence of DNA2-i (Fig. 6A) then performed 2D gel analysis (as in Fig. 3A-C) to monitor the signal from double-Ys and reversed forks (Fig. 6B-D). In the vehicle control, double-Ys increased in mobility over time (Fig. 6B,i-iv) indicating that they became more extensively degraded. This was accompanied by loss of double-Y signal (Fig. 6C), as previously observed (Fig. 3D). DNA2-i treatment lessened the decrease in mobility of double-Ys (Fig. 6B,iii,vi) and increased the abundance of double-Ys (Fig. 6C, Supplemental Fig. S7B), as expected if degradation can initiate at replication forks (Supplemental Fig. S7A,iv). These data cannot be explained by degradation and then restoration of reversed forks because double-Ys were progressively converted to reversed forks (Fig. S7B-C). Thus, our data support a model where degradation can occur prior to fork reversal.

**Figure 6:**
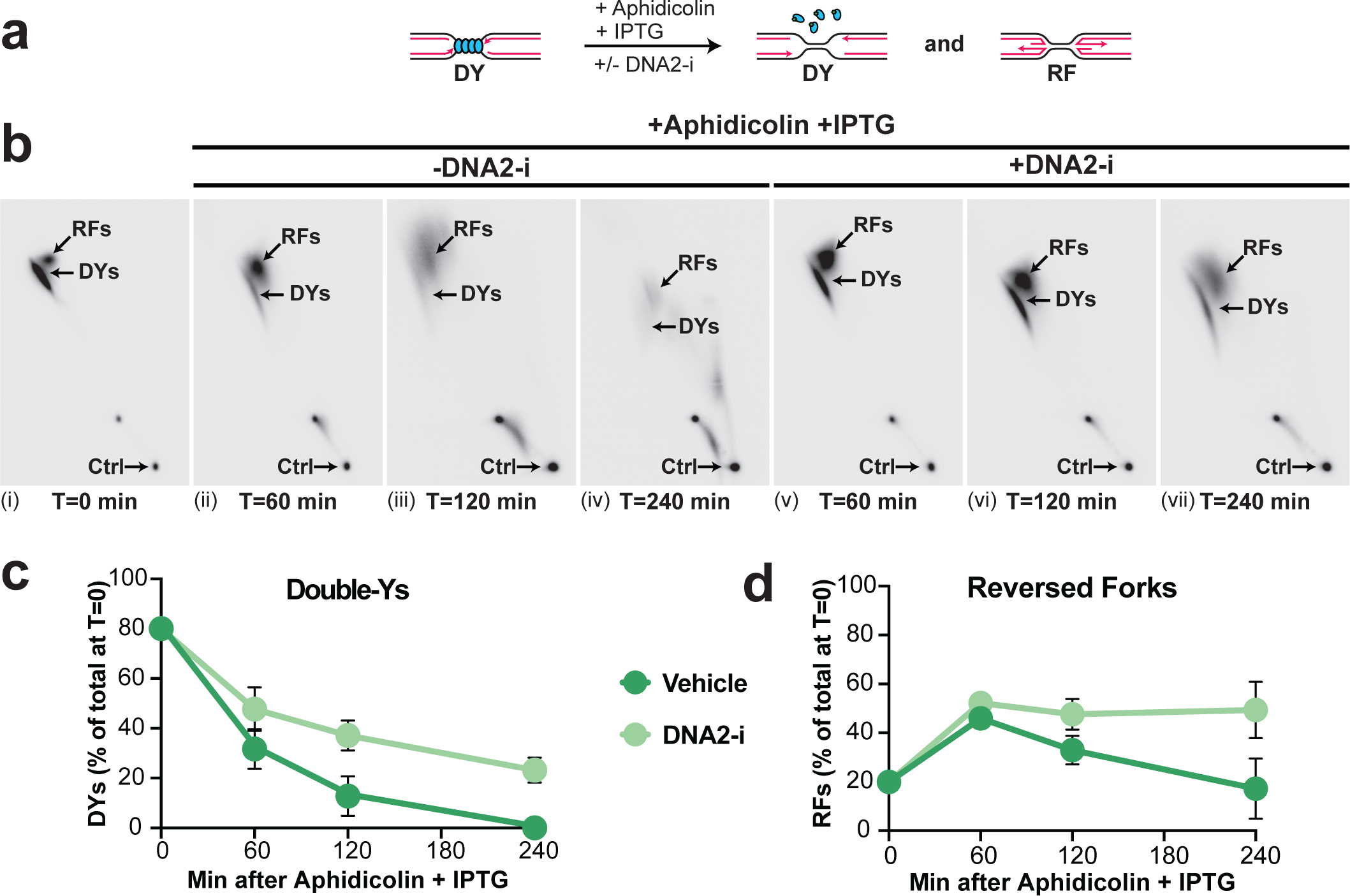
DNA2 degrades replication forks and reversed forks. *(A)* Forks were localized to a LacR barrier and NSD was induced by addition of IPTG and aphidicolin in the absence or presence of DNA2-i. Purified DNA was subjected to restriction digest so that replication fork structures could be visualized as DYs. *(B)* Double-Y (DY) and reversed fork (RF) structures from (a) were separated by 2-D gel electrophoresis (as in Fig. 3B-C). *(C)* Quantification of DY structures from (b) as a percentage of total signal at T=0. Mean ± S.D., n=3 independent experiments. See also Supplemental Fig. S7B. *(D)* Quantification of RF structures from (d) as a percentage of total signal at T=0. See also Supplemental Fig. S7C.

Our time course 2D gel analysis revealed two further observations that further supported our model that Y-shaped forks are degraded prior to reversed forks. First, most double-Ys increased in mobility by 60 minutes (Fig. 6B,i), indicating that they underwent significant degradation by this time point. Importantly, reversed fork degradation took place later, from 60 to 240 minutes (Fig. 6D). Thus, double-Y degradation took place prior to reversed fork degradation. Additionally, DNA2-i treatment essentially blocked degradation of reversed forks during our experiments (Fig. 6D), indicating that it was required for degradation of reversed forks. However, in the presence of DNA2-i double-Ys still increased in mobility (Fig. 6B,v) and decreased in abundance (Fig. 6C). Thus, double-Y degradation took place even when reversed forks were not degraded. These observations definitively show that NSD at Y-shaped forks can take place prior to, and independent of, degradation at reversed forks.

## Discussion

We developed an *in vitro* approach to study nascent strand degradation in vertebrates. We have demonstrated that NSD of both strands is an initial response to fork stalling and involves progressive conversion of uncoupled forks to reversed forks. Importantly, NSD and fork reversal are stimulated by replication fork uncoupling, rather than fork stalling (Fig. 7A-B). Additionally, NSD occurs at replication forks prior to fork reversal indicating that an extra degradation step (Fig. 7B-C) exists, in addition to the degradation previously reported at reversed forks (Fig. 7D-E). The replicative CMG helicase remains associated with DNA during NSD and fork reversal, suggesting that it continues to encircle ssDNA during fork reversal and nascent strand degradation (Fig. 7C-D). These observations inform a new model for NSD (Fig. 7) and have implications for fork reversal and template switching (Supplemental Fig. S7D). These points are discussed further below.

**Figure 7:**
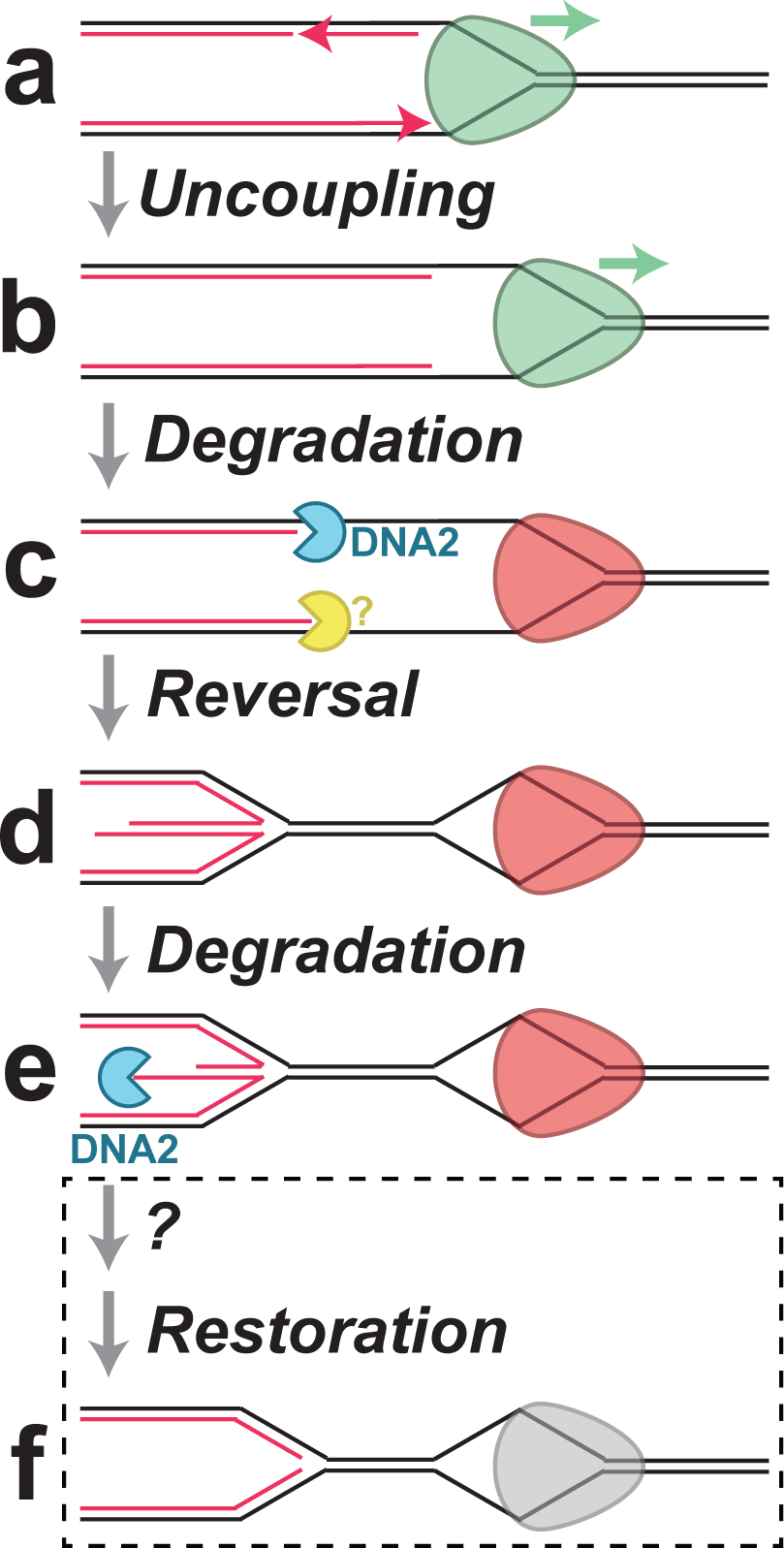
Model for nascent strand degradation in *Xenopus* egg extracts. *(A)* Replicative helicase and DNA polymerase activities are coupled at replication forks. *(B)* When DNA polymerases stall the replicative helicase continues unwinding, resulting in uncoupling. *(C)* Uncoupling initially stimulates NSD by DNA2 and one or more other nucleases. The uncoupled replicative helicase stalls (red). *(D)* Uncoupling causes replication fork reversal. The replicative helicase remains bound to single-stranded DNA suggesting parental DNA strands are reannealed behind it. *(E)* The regressed arm undergoes NSD, which requires DNA2. *(F)* Restoration of the reversed arm is likely to involve an additional step that has not been identified and it is unclear whether the replicative helicase would be active at this point.

### Consequences of CMG helicase retention

Replication fork DNA protects the CMG helicase from the replisome removal pathways that operate during replication termination^50, 51^. Our observation that CMG is retained on DNA (Fig. 5C) during NSD and fork reversal (Fig. 4C,F) means that CMG must remain at a replication fork structure during these processes. The simplest explanation is that parental DNA is reannealed in the wake of CMG, which would then reside within a single-stranded DNA bubble (Fig. 7C-D). This model is appealing because it would allow CMG to readily restart DNA synthesis following restoration of a reversed fork structure and extension of the nascent DNA strands. It would also limit generation of ssDNA and thus prevent depletion of RPA, which is toxic to cells^22^. Additionally, reannealing of nascent DNA strands would create a second replication fork structure devoid of replication proteins, which may favor the activity of fork reversal enzymes which would otherwise need to compete with CMG for replication fork DNA^7, 30, 52^. However, such a mechanism would require ssDNA helicases to unwind parental DNA between the nascent strands and the CMG helicase following fork restoration (Supplemental Fig. S7D,v). It will be important to address whether any of the DNA helicases implicated in fork restart^1–4^ or counteracting fork reversal^53^ fulfill this role.

### Implications for replication restart

We observed that uncoupled replication forks were progressively converted to reversed forks following uncoupling (Fig. 3E). This is surprising because replication forks and reversed forks are thought to undergo interconversion^1, 6, 7^. Because reversed forks need to be converted to replication forks to resume DNA synthesis some sort of restoration event is needed to convert reversed forks to replication forks (Fig. 7E-G). Since our experiments block DNA-synthesis it likely that restoration of reversed forks involves DNA synthesis. This would be consistent with the reported antagonism between repriming and fork reversal pathways^54, 55^. It will be crucial to determine which other processes cooperate with NSD and fork reversal enzymes to convert reversed forks back to replication forks.

### Multiple nascent strand degradation events

Our data show that DNA2 degrades uncoupled forks prior to fork reversal (Fig. 6, Fig. 7C-D). Previous work in yeast suggested that DNA2 activity at forks might inhibit fork reversal^56^. While we observed increased reversed fork signal following inhibition of DNA2 (Fig. 6D), we did not observe increased frequency of reversed fork formation, arguing against this role for DNA2 in our experiments (Supplemental Fig. S7C). Instead, our results support a role for DNA2 in degradation reversed forks (Fig. 6D), as previously reported^8^.

It is unclear what function is fulfilled by degradation of uncoupled forks. DNA2 promotes extensive NSD in at least two distinct fork reversal pathways^38^, which would be consistent with a role upstream of fork reversal. Given its role in degrading reversed forks^8^, we favor the idea that DNA2 activity at uncoupled forks somehow promotes degradation of reversed forks that are subsequently formed. For example, DNA2 might degrade uncoupled forks to ensure that the 5’ strand of the subsequent reversed fork is recessed and primed for extensive degradation^45^. It will be important to establish the role of NSD at uncoupled forks and whether it is negatively regulated by any fork protection proteins in the same way that NSD at reversed forks is^9, 10, 13^.

### How does uncoupling elicit fork reversal and NSD?

The observation that uncoupling stimulates both nascent strand degradation and fork reversal (Fig.4, Fig. 7A-C) is consistent with a correlation between uncoupling and replication fork reversal^14^. However, it is unclear exactly which molecular event involved in uncoupling causes NSD and fork reversal. One possibility is that nascent strands efficiently stimulate fork reversal and NSD but this is typically inhibited by interactions between the replisome and the nascent DNA strands. In this view uncoupling would physically separate the replisome from the nascent strands to grant access to the fork processing and remodeling enzymes. Alternatively, the ssDNA generated by CMG helicase activity during uncoupling may stimulate reannealing of the parental strands and generate the substrate for fork reversal. This would be consistent with other situations where NSD and processing occur at post-replicative gaps^10, 55, 57^. It will be important to address whether any of the proteins that modulate post-replicative gap stability independent of fork reversal^58, 59^ are also involved in limiting the initial NSD step we have identified at forks (Fig. 7b-c).

Although uncoupling is clearly important for both NSD and fork reversal, our experiments did not demonstrate an absolute requirement for uncoupling in either process. NSD was detected even when uncoupling was inhibited (Fig. 4C) and fork reversal was detected prior to uncoupling (Fig. 3E). These observations may reflect our inability to completely block uncoupling (Fig. 4C) or low levels of uncoupling during replication (Fig. 3E). Alternatively, there may be other triggers for both fork reversal and NSD that can operate in different contexts, consistent with multiple pathways for NSD and fork reversal^38^. It will be crucial to identify the full set of triggers and pathways for NSD and fork reversal.

## Methods

### *Xenopus* egg extracts

*Xenopus* egg extracts were prepared from *Xenopus laevis* wild-type males and females (Nasco) as previously described^60^ and approved by Vanderbilt Division of Animal Care (DAC) and Institutional Animal Care and Use committee (IACUC).

### Plasmid Construction and Preparation

pJD145, pJD156, and pJD90 were described previously^41^. To create pJD161 (p[*lacO*x25]-XhoI-[*lacO*x25]) DNA oligonucleotides JDO120 (GTACAAGTAAATCAGAGCCAGATTTTTCCTCCTCTCGGAATTGTGAGCGGATAACAATTCCCTCGAGCCAATTGTGAGCGGATAACAATTGGAAGTGCAGAACCAATGCATGCAGGAGATTTGAC) and JDO121 (GT ACGTCAAATCTCCTGCATGCATTGGTTCTGCACTTCCAATTGTTATCCGCTCACAATTGGCTCGAGGGAATTGTTATCCGCTCACAATTCCGAGAGGAGGAAAAATCTGGCTCTGATTTACTT) were annealed to create a DNA fragment that contained a XhoI site flanked by a lacO sequence on each side. The JDO120/JDO121 duplex was then cloned into the BsrGI site of pJD90 (p[*lacO*x24]). This cloning procedure regenerated the BsrGI site, which was then used as the target for insertion of a BsrGI-BsiWI fragment from pJD90 that contained 24 tandem *lacO* repeats. The resulting plasmid was then digested with BsrGI and BsiWI to create a fragment containing 50 tandem *lacO* repeats with a XhoI site in the middle. This fragment was inserted into the BsiWI site of pJD145 to yield pJD161.

### DNA replication in Xenopus egg extracts

High Speed Supernatant (HSS) was supplemented with nocodazole (3 ng/μl) and ATP regenerating system (ARS; 20mM phosphocreatine, 2 mM ATP and 5 ng/μl creatine phosphokinase) then incubated at room temperature for 5 minutes. To license plasmid DNA, 1 volume of ‘licensing mix’ was prepared by adding plasmid DNA to HSS at a final concentration of 15 ng/μl, followed by incubation at room temperature for 30 minutes. NucleoPlasmid Extract (NPE), extract was supplemented with ARS, DTT (final concentration: 2 mM), [α-^32^P]dATP (final concentration: 350 nM) and diluted to 45 in 1X Egg Lysis Buffer (ELB, 250 mM Sucrose, 2.5 mM MgCl2, 50 mM KCl, 10 mM HEPES, pH 7.7). To form a replication barrier, LacR was bound to *lacO* repeats as previously described^41^. To initiate replication, 2 volumes of NPE mix were added to 1 volume of Licensing mix. Replication forks were stalled at the LacR-bound *lacO* array and then released by IPTG addition as described previously. Reactions were stopped by addition of 10 volumes Extraction Stop Solution (0.5 SDS, 25 mM EDTA, 50 mM Tris-HCl, pH 7.5). Samples were subsequently treated with RNase A (final concentration: 190 ng/µl) and then Proteinase K (909 ng/µl) before either direct analysis by gel electrophoresis or purification of DNA as described previously^41^.

For most experiments pJD156 (p[*lacO*x32]) was used as the template for replication. For the experiments in Fig. 3F-H and Supplemental Fig. S4J-K pJD161 (p[*lacO*x25]-XhoI-[*lacO*x25]) was used. pJD145 (p[CTRL]) did not contain a *lacO* array, which allowed it to fully replicate in the presence of LacR, and was used as a loading control because its small size allowed it to be readily distinguished from pJD156 and pJD161. In most experiments, pJD145 was added to the licensing mix at a final concentration of 1 ng/μl and the plasmid replicated prior to induction of NSD due to the absence of any LacR array. In Supplemental Fig. S1C-I radiolabeled pJD145 was added to NPE at a concentration of 1 ng/μl prior to initiation of DNA replication and the plasmid did not replicate because it had not undergone prior licensing of the DNA.

Aphidicolin from *Nigrospora sphaerica* (Sigma-Aldrich) was dissolved in DMSO and added to reactions at a final concentration of 330 μM. Mirin (Selleckchem) was dissolved in DMSO and added to reactions at a final concentration of 500 μM. C5 (AOBIOUS) was dissolved in DMSO and added to reactions at a final concentration of 3.5 mM. NMS-873 (Selleckchem) was dissolved in DMSO and added to reactions at a final concentration of 200 μM. For drugs dissolved in DMSO the final reaction concentration of DMSO was 4% (V/V).

### Protein Purification

Biotinylated LacR was expressed in *Escherichia coli* and purified as described previously^41^.

### Antibodies

Antibodies against Cdc45, Mcm6 and RPA were previously described^42^.

### Nascent Strand Degradation assays

To monitor DNA synthesis, samples were separated on a 1% agarose gel at 5 V/cm. Radiolabeled DNA was visualized by phosphorimaging to measure incorporation of radiolabeled nucleotides. Signal was quantified using ImageQuant (GE Healthcare) and ImageJ and normalized to the loading control in each lane.

To monitor NSD, samples were purified and then digested with 0.4 U/μl XmnI in CutSmart Buffer (NEB) for 30 minutes at 37°C and then separated on a 1% agarose gel at 5 V/cm. Radiolabeled DNA was detected by phosphorimaging and Double-Y signal was quantified and normalized to pJD145 signal, which served as a loading control.

To monitor disappearance of intact nascent strands, purified NSD intermediates were digested with 0.4 U/μl AlwnI (NEB) in CutSmart Buffer (NEB) for 1 hour at 37°C. Digest was stopped by adding EDTA to a final concentration of 30 mM. Reaction was then prepared for electrophoresis by adding Alkaline Loading Buffer 6X (EDTA, Ficoll, Bromocresol green, Xylene Cyanol and NaOH (10N)) to a final concentration of 1X. Nascent strands were then separated on a 1.5% denaturing alkaline gel at 1.5 V/cm. Denaturing gel was then neutralized by gentle agitation in 7% TCA solution and radiolabeled DNA was detected by phosphorimaging as described above.

### 2D gel electrophoresis

2D gels were performed as described ^42^ with slight modifications. Briefly, purified DNA was digested with XhoI and DraIII (Fig. 4H) or XmnI (all other gels) and digested DNA was separated on a 0.4% agarose gel at 1 V/cm for 22 hours. A second dimension gel containing 1.2% agarose and 0.3 µg/ml ethidium bromide was cast over the first dimension gel slice and separated at 5 V/cm for 12 hours at 4°C. Radiolabeled DNA was then detected by phosphorimaging as described above.

### Nick translation of DNA

To radiolabel parental DNA strands, 60 ng/µl plasmid DNA was resuspended in 1X NEB Buffer 2.1 and treated with 0.5 U/µl Nt.BbvCI at 37°C for 1 hour, followed by heat inactivation at 80°C for 20 minutes. To perform nick translation the reaction was supplement with 0.5 volumes of dNTPs (12.5 mM each of dCTP, dGTP, dTTP), 3 U/µl *E. coli* DNA Polymerase 1, and 0.33 mM [α^32^P]-dATP in 1X NEB Buffer 2.1 and incubated at 16°C for 10 minutes then on ice for 5 minutes. Nick-translated DNA was purified into 10 mM Tris-HCL, pH 8.0 using micro Bio-Spin columns (Bio-Rad) and used for replication and NSD assays.

### Plasmid pull downs

Plasmid pull downs were performed as previously described^42^.

## Acknowledgements

JMD was supported by NIH grant R35GM128696, ACS grant IRG-15-169-56, and Vanderbilt-Ingram Cancer Center Support Grant (P30CA068485).

## Supplemental Figure Legends

**Supplemental Fig. S1.**
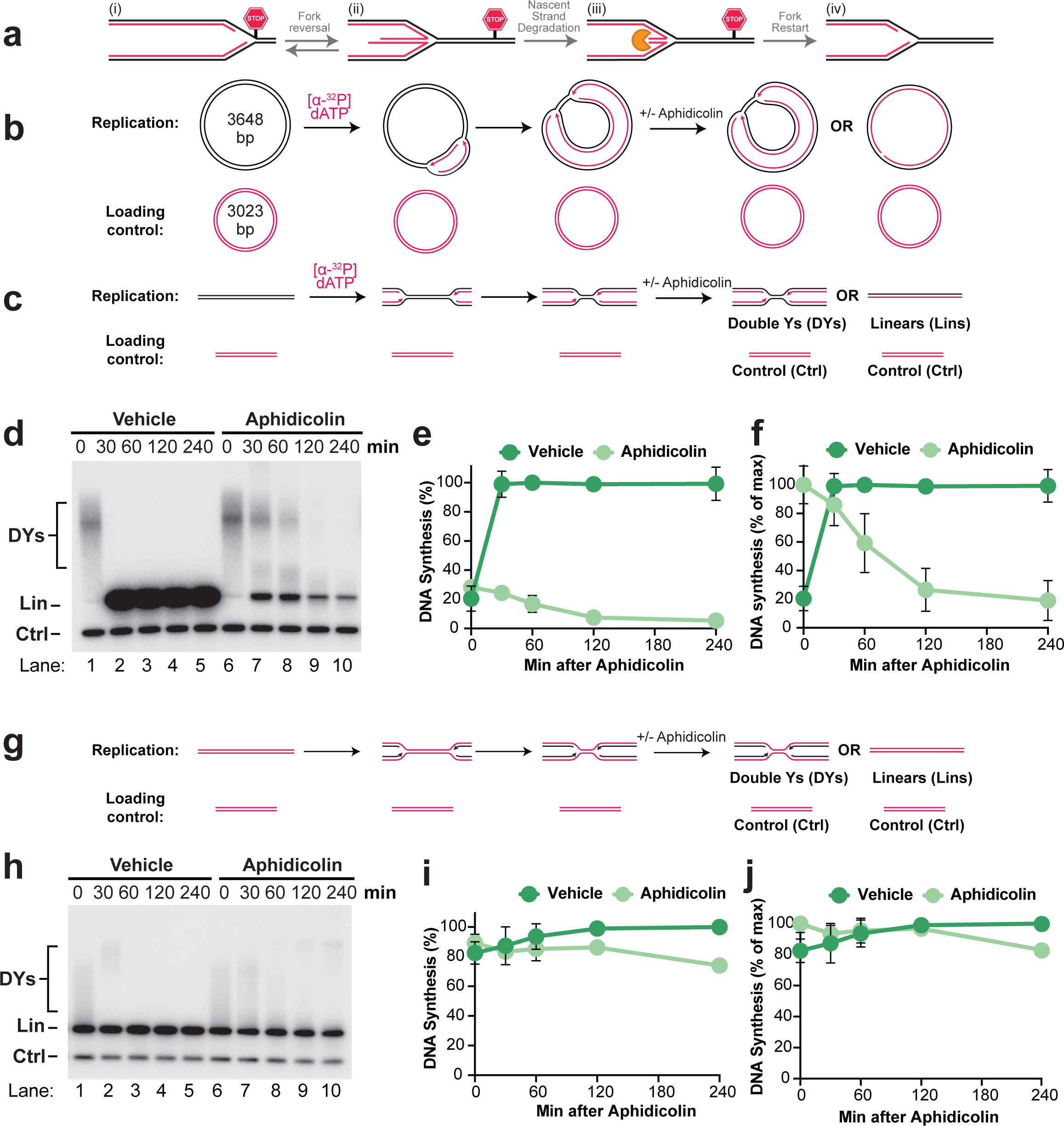
Nascent strand degradation is an initial response to fork stalling. **A)** A current model for fork reversal and nascent strand degradation. **B)** Plasmid DNA was replicated using *Xenopus* egg extracts and newly-synthesized nascent strands were radiolabeled by inclusion of [α-^32^P]dATP. 6 minutes after initiation, reactions were treated with aphidicolin, to inhibit DNA synthesis, or vehicle control. As a loading control (Ctrl) the reactions include a smaller plasmid that was radiolabeled prior to the experiment and did not undergo replication. **C)** DNA structures from (b) were purified and digested with XmnI, which cuts the plasmid once. **D)** Samples from (c) were separated on an agarose gel and visualized by autoradiography. **E)** Quantification of DNA synthesis from (d) normalized to the maximum signal across all time points and conditions. Mean ± S.D., n=3 independent experiments. **F)** Quantification of DNA synthesis in (d) normalized to the maximum signal for each condition across all time points. Mean ± S.D., n=3 independent experiments. **G)** Pre-radiolabeled plasmid DNA was replicated by addition of *Xenopus* Nucleoplasmic Extract (NPE). 6 minutes after initiation, DNA synthesis was inhibited by the addition of Aphidicolin. **H)** XmnI digested molecules were separated on an agarose gel and visualized by autoradiography. As a loading control (Ctrl) the reactions include a smaller radiolabeled plasmid that did not undergo replication. **I)** Quantification of (h) as in (d). Mean ± S.D., n=3 independent experiments. **J)** Quantification of (h) as in (e). Mean ± S.D., n=3 independent experiments.

**Supplemental Fig. S2.**
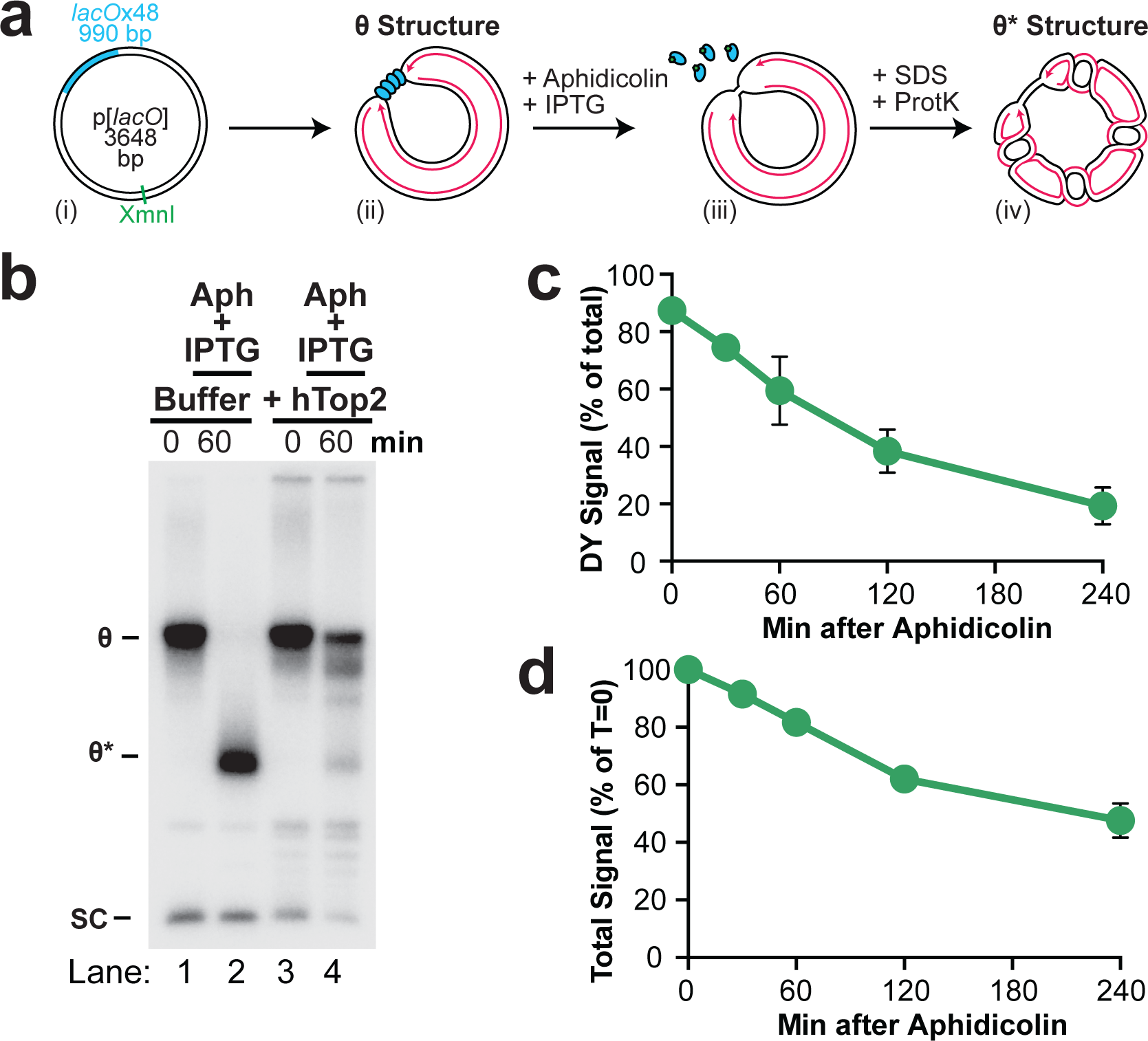
Characterization of localized, synchronous nascent strand degradation. **A)** Cartoon depicting the source of θ* structures in Fig. 1B. The XmnI site is located 1013 base pairs away from the closest edge of the *lacO* array. **B)** DNA structures from Fig. 1A were treated with human topoisomerase II (hTop2) or buffer control, then separated on an agarose gel and visualized by autoradiography. hTop2 treatment converted θ* signal back to θs, demonstrating that θ* structures are topoisomers of θs. **C)** Quantification of DYs as a % of total lane signal from Fig. 1E. Mean ± S.D., n=5 independent experiments. **D)** Quantification of total lane signal from Fig. 1E. Mean ± S.D., n=5 independent experiments.

**Supplemental Fig. S3.**
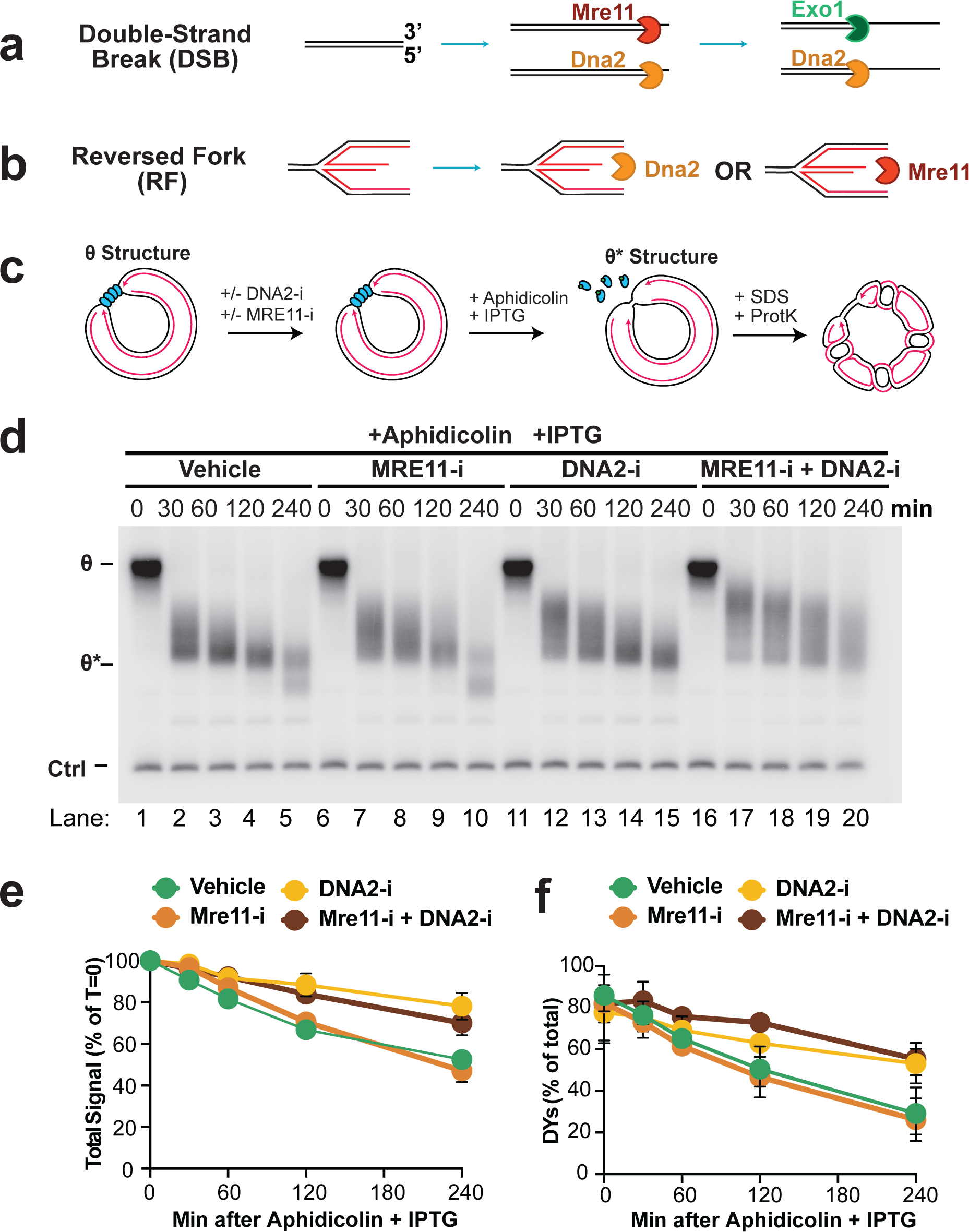
Characterization of MRE11 and DNA2 activity during nascent strand degradation. **A)** Cartoon depicting the roles of MRE11 and DNA2 in resection at double-strand breaks **B)** Cartoon depicting the roles of MRE11 and DNA2 in resection at reversed forks **C)** Depiction of the DNA structures generated by Fig. 2D, prior to restriction digest. **D)** DNA intermediates from (c) were separated on an agarose gel and visualized by autoradiography. **E)** Quantification of DY signal in Fig. 2E as a percentage of total signal for each lane. Mean ± S.D., n=3 independent experiments. **F)** Quantification of total lane signal from Fig. 2E. Mean ± S.D., n=3 independent experiments.

**Supplemental Fig. S4.**
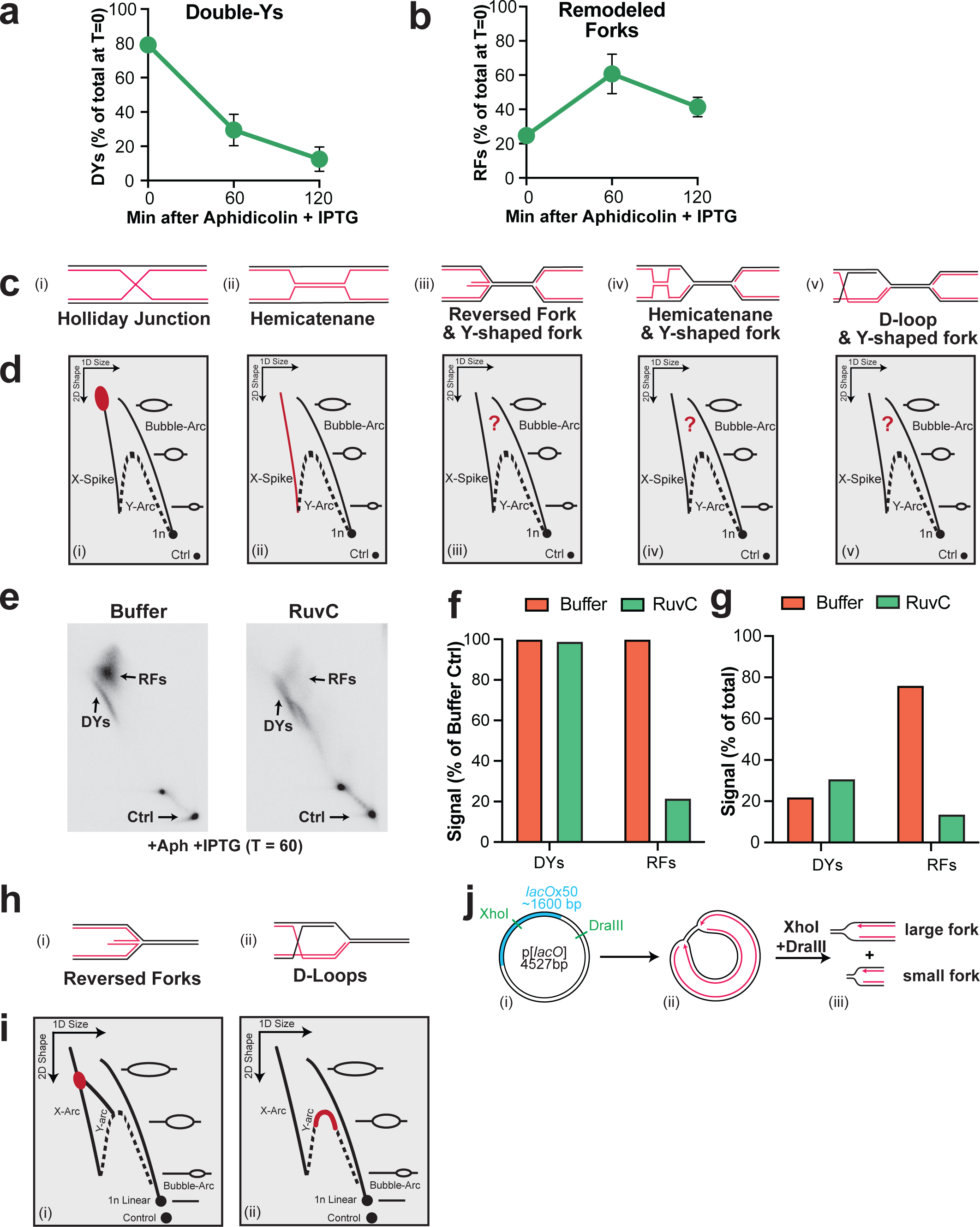
Characterization of DNA structures detected by 2-D gel during nascent strand degradation. **A)** Quantification of Double-Ys from Fig. 3C expressed as a percentage of total signal at T=0. Mean ± S.D., n=3 independent experiments. **B)** Quantification of remodeled forks from Fig. 3C expressed as a percentage of total signal at T=0. Mean ± S.D., n=3 independent experiments. **C)** Cartoon indicating the different DNA structures that the remodeled forks in Fig. 3C could correspond to. **D)** Expected 2-D gel migration pattern of the structures depicted in (c). The region highlighted in red indicates where each structure is expected to migrate. ‘?’ indicates structures whose expected migration on a 2-D gel is unclear. **E)** Samples from Fig. 3C,iii were treated with buffer or RuvC, separated by 2-D gel electrophoresis and visualized by autoradiography. **F)** Quantification of DYs and RFs following RuvC treatment in (e) expressed relative to the abundance of each structure in the buffer treated condition. **G)** Quantification of RFs and DYs from (c) expressed relative to total signal on each gel. **H)** Cartoon indicating the structure of a D-loop and a reversed fork. **I)** Expected 2-D gel migration pattern of the structures depicted in (h). The region highlighted in red indicates where each structure is expected to migrate. **J)** Cartoon indicating the restriction digest strategy used to excise individual replication forks for 2-D gel analysis. The DraIII site is 450 bp away from the nearest edge of the *lacO* array and 1250 bp away from the XhoI site.

**Supplemental Fig. S5.**
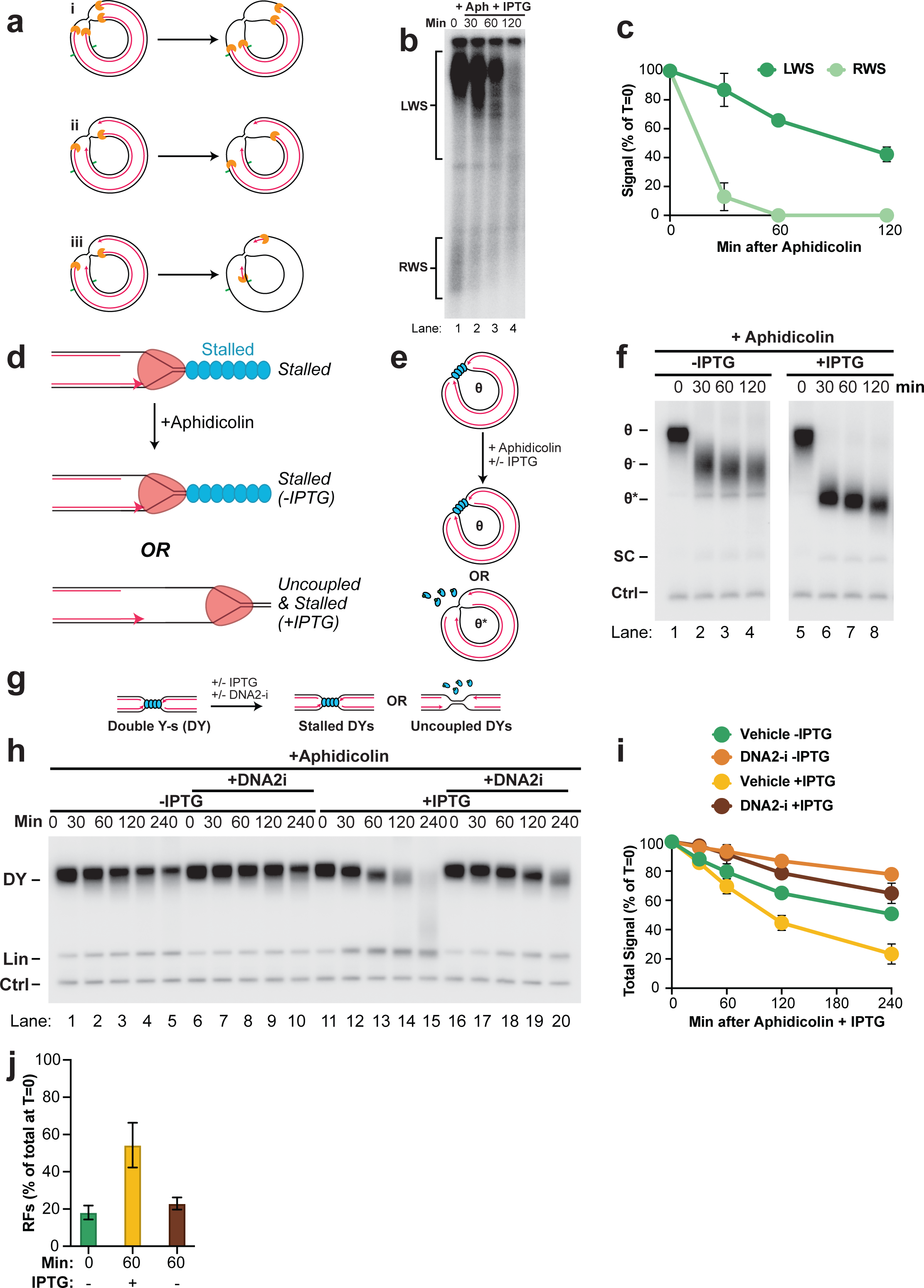
Characterization of stalled and uncoupled replication forks. **A)** Depiction of different models for degradation of leading and lagging strands during NSD observed in Fig. 4 and the expected effect on degradation of LWS and RWS depicted in Fig. 4A. In (i) both leading and lagging strand are degraded simultaneously and synchronously, which should result in disappearance of RWS before LWS. In (ii) lagging strands only are degraded synchronously, which should result in persistence of ∼50% of RWS and LWS until the leading strand are degraded by nuclease activity that initiated at lagging strands of the opposite fork. This would result in biphasic kinetics of degradation. In (iii) lagging strands only are degraded asynchronously, which should result in loss of RWS and LWS at the same rate. **B)** Uncropped gel of Fig. 4B lanes 1-4 shown after a longer exposure so that the weaker rightward strands (RWS) could be visualized. **C)** Quantification of LWS and RWS from (b). RWS are degraded before LWS, with no evidence of biphasic kinetics, indicating that both strands are degraded simultaneously, as in (a,i). Mean ± S.D., n=3 independent experiments. **D)** Cartoon depicting the strategy to test the role of helicase stalling during NSD. If helicase stalling triggers NSD then NSD should be increased in the absence of IPTG. If uncoupling triggers NSD then addition of IPTG should increase NSD. **E)** Depiction of the DNA structures generated by Fig. 4A, prior to restriction digest. **F)** Samples from (e) were separated on an agarose gel and visualized by autoradiography. θ^-^ species arise from the low level of uncoupling that occurs even in the presence of LacR, which is not a complete block to helicase progression. **G)** NSD was induced as in Fig. 4A in the presence or absence of DNA2-i. **H)** Samples from (g) were separated on a native agarose gel and visualized by autoradiography. **I)** Quantification of DY signal from (h). Mean ± S.D., n=3 independent experiments. **J)** Quantification of reversed forks from Fig. 4E expressed as a percentage of total signal at T=0. Mean ± S.D., n=3 independent experiments.

**Supplemental Fig. S6.**
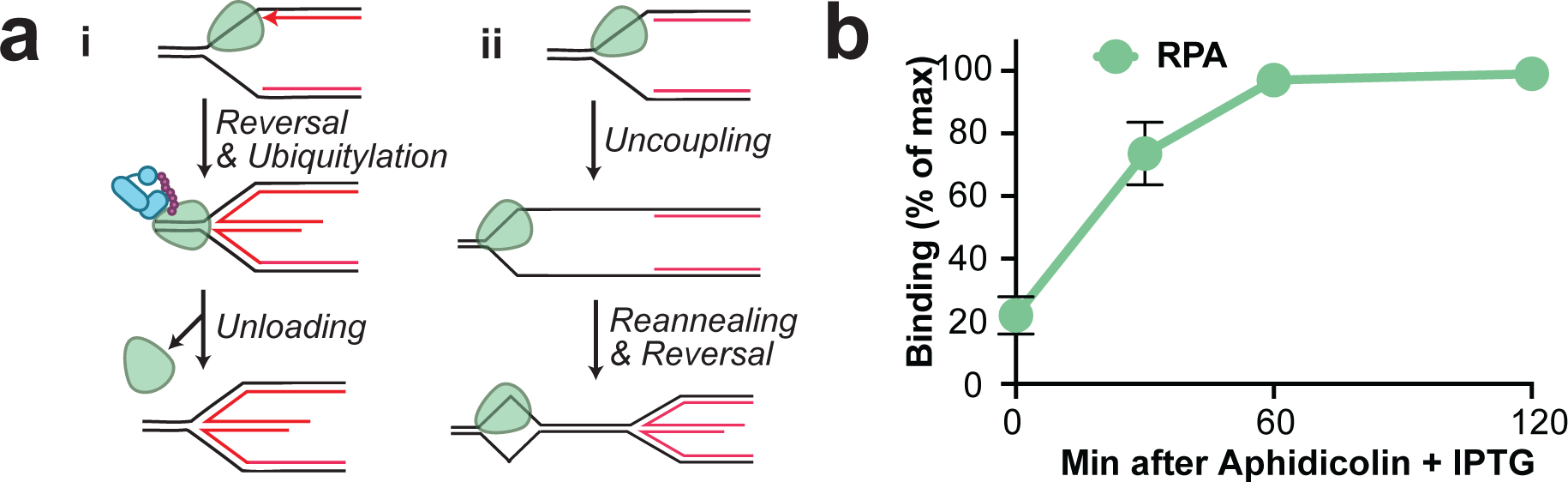
Binding of replication fork proteins during NSD. **A)** Cartoon of two different models of CMG helicase behavior during fork reversal and NSD. In (i) the CMG helicase translocates onto double-stranded DNA, as suggested, and then is removed from DNA, as previously proposed^1^. In (ii) the replisome remains on DNA, suggesting that it resides in a ssDNA bubble ahead of the reversed fork. **B)** Quantification of RPA signal from Fig. 5B. Mean ± S.D., n=3 independent experiments.

**Supplemental Fig. S7.**
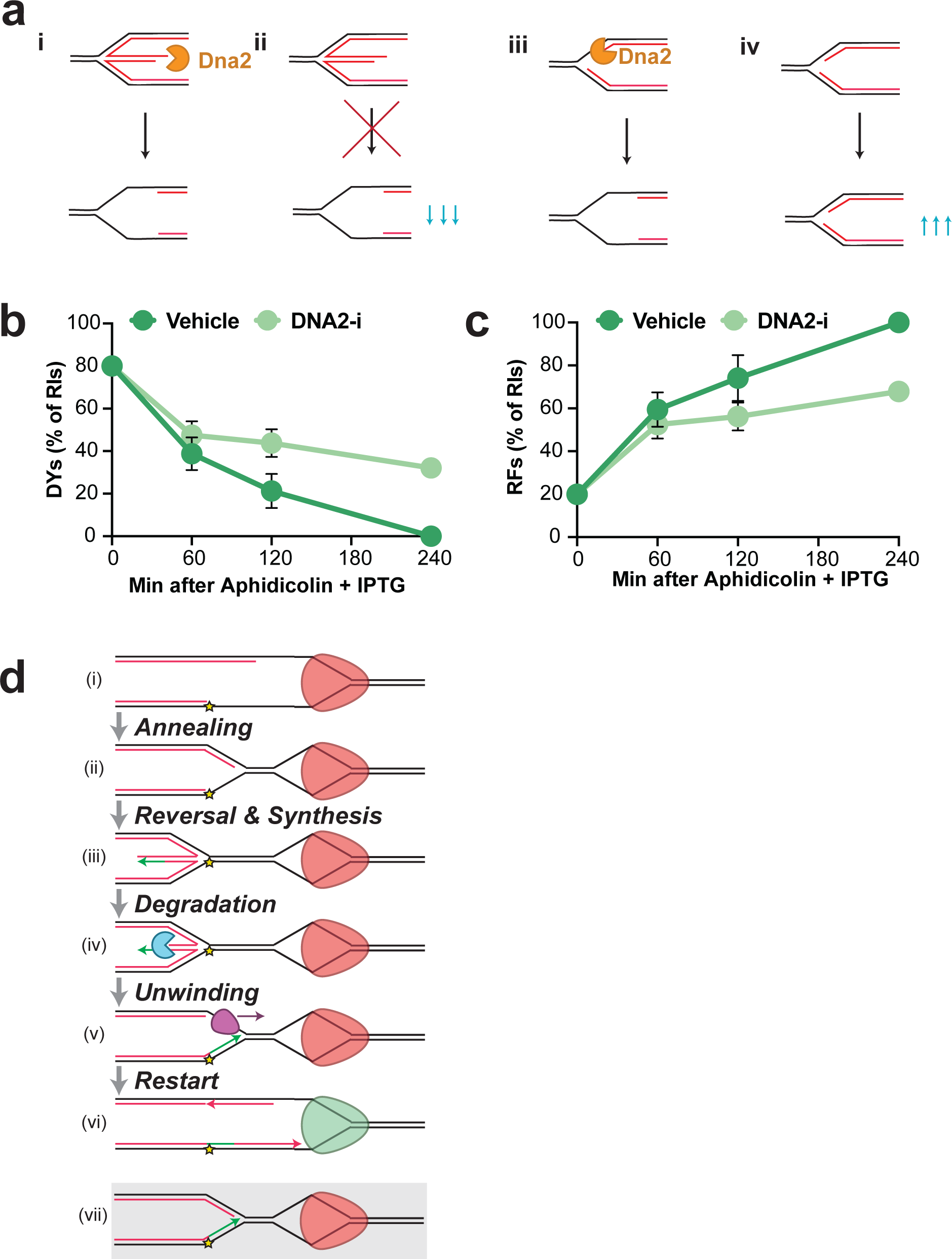
Additional degradation steps during NSD. **A)** Cartoon of different models to explain a role for DNA2 in degrading Y-shaped forks during NSD and the expected impact of loss of DNA2 activity on Y-shaped forks in each case. (i) depicts a model where DNA2 degrades reversed forks to generate Y-shaped forks. In this model the Y-shaped forks are degraded as consequence of prior DNA2 activity at reversed forks. (ii) depicts the impact of impaired DNA2 activity on (i). Y-shaped forks should decrease in abundance as less degradation of reversed forks should reduce the formation of Y-shaped forks. (iii) depicts a model where DNA2 degrades forks prior to fork reversal. (iv) depicts the impact of impaired DNA2 activity on (iii). Y-shaped forks should increase in abundance due to less degradation of nascent strands, which is the source of the radioactive signal that is measured. **B)** Quantification of Double-Ys from Fig. 5B expressed as a percentage of total replication intermediates (RIs). Mean ± S.D., n=3 independent experiments. **C)** Quantifications of reversed forks from Fig. 5B expressed as a percentage of total replication intermediates (RIs). Mean ± S.D., n=3 independent experiments. **D)** Potential model for template switching, based on our new model for NSD (Fig. 7). (i) after encountering a polymerase-blocking lesion (yellow) replication forks uncouple, resulting in a stalled replisome (red). (ii) The replisome is retained on DNA and parental DNA strands anneal to create a second replication fork that is the substrate for fork reversal. This places the replisome within a single-stranded DNA bubble. (iii) The nascent lagging strands serve as template for leading strand synthesis (green). (iv) Exonuclease activity at the extruded DNA end generates a 3’ overhang. (v) restoration of the reversed fork results in a lagging strand gap that allows loading of a 5’-3’ single-stranded DNA helicase that can unwind the re-annealed parental strands. (vi) nascent strands are extended to the replisome, which resumes unwinding (green). (vii) Without exonuclease activity at the extruded DNA end (in (iv)) restoration of the reversed fork would result in a 3-way fully duplex fork structure that could not be readily unwound.

